# A structural basis for prion strain diversity

**DOI:** 10.1101/2022.05.17.492259

**Authors:** Szymon W. Manka, Adam Wenborn, Jemma Betts, Susan Joiner, Helen R. Saibil, John Collinge, Jonathan D.F. Wadsworth

## Abstract

Recent cryo-EM studies of infectious, ex vivo, prion fibrils from hamster 263K and mouse RML prion strains revealed a comparable, parallel in-register intermolecular β-sheet (PIRIBS) amyloid architecture. Rungs of the fibrils are composed of individual prion protein (PrP) monomers that fold to create distinct N- and C-terminal lobes. However, disparity in the hamster/mouse PrP sequence precludes understanding how divergent prion strains emerge from an identical PrP substrate. Here, we determined the near-atomic resolution cryo-EM structure of infectious, *ex vivo* mouse prion fibrils from the ME7 prion strain and have compared this with the RML fibril structure. This structural comparison of two biologically distinct mouse-adapted prion strains suggests defined folding sub-domains of PrP rungs and the way in which they are interrelated, providing the first structural definition of intra-species prion strain-specific conformations.

## Introduction

Prions are infectious agents causing invariably fatal neurodegenerative diseases in mammals including Creutzfeldt-Jakob disease (CJD) in humans, scrapie in sheep and goats, bovine spongiform encephalopathy (BSE) in cattle and chronic wasting disease (CWD) in cervids^1-3^. Prions are composed of polymeric fibrillar assemblies of misfolded host-encoded cellular PrP (PrP^C^) which propagate by fibre elongation and fission. Biologically distinct prion strains can however be serially propagated in identical hosts expressing the same PrP^C^ and produce different disease phenotypes. Understanding the structural basis of prion strain diversity is of considerable biological interest and evolutionary significance. In addition, prions may transmit disease between mammalian species, as with the infection of humans with BSE prions causing variant CJD. Such cross-species transmission is limited by so-called species barrier effects which relate to structural compatibility of prion strains with host PrP^C^ according to the conformational selection model^2,4^. As it is well recognised that novel prion strains with altered host ranges can arise as a result of PrP polymorphisms in both inter- and intra-species transmissions^2,3,5^, determining the structural basis of prion diversity is critical to understanding whether emerging animal prion strains constitute a zoonotic risk to public health. Furthermore, accumulating evidence suggests that analogous mechanisms of seeded polymerisation and spread of host proteins is involved in the pathogenesis of multiple neurodegenerative conditions, notably Alzheimer’s and Parkinson’s diseases, where distinct strains of protein assemblies have also been postulated^3,6-9^.

Although it is now firmly established that mammalian prions are composed of fibrillar assemblies of misfolded host-encoded PrP, and propagate by means of seeded protein polymerization and fission^1-3,10-16^, the structural mechanisms underpinning prion strain diversity remain unclear. While it is known that prion strains represent distinct misfolded PrP conformations and assembly states^1-3,17-21^, high-resolution structural definition of prion strains has been extremely problematic. In this regard, because *in vitro*-synthetically generated PrP amyloids are either devoid of detectable prion infectivity or have specific-infectivities too low for meaningful structural analysis^2,3,13,22^, efforts to define authentic infectious prion structures have to overcome the difficulty of isolating relatively homogeneous *ex vivo* prion strain assemblies of correspondingly extremely high-specific infectivity suitable for structural analysis^13,23^. Recently however, significant progress has been made^14-16^.

High-resolution cryo-EM studies of infectious, *ex vivo*, prions isolated from the hamster 263K prion strain^14^ or the mouse RML prion strain^15^ reported single-protofilament helical amyloid fibrils that have a broadly similar, parallel in-register intermolecular β-sheet (PIRIBS) amyloid architecture. Rungs of the fibrils are composed of single PrP monomers that fold to create distinct N- and C-terminal lobes with the N-linked glycans and glycosylphosphatidylinositol (GPI) anchor projecting from the C-terminal lobe^14,15^. Transgenic mice expressing GPI-anchorless PrP when infected with RML prions generated in wild-type mice propagate prion fibrils with the same fold as seen in RML prion-infected wild-type mice^15,16^. The overall architecture of hamster 263K and mouse RML fibrils are remarkably similar and compatible with the defining physicochemical properties of prions^14-16^.

Despite the overall similarity of the hamster 263K and mouse RML prion fibril architectures, there are pronounced differences in the fold of the C-terminal lobes^15,16^, which may be attributable to differences in PrP amino acid sequence and/or distinct conformations associated with divergent prion strains. To determine directly which conformational differences can be attributed to prion strain, comparison of structures of different strains propagated in the same host with identical PrP^C^ substrate is necessary.

Here we present a 2.6 Å cryo-EM structure of fibrils present in highly infectious prion rod preparations isolated from the brain of mice infected with the ME7 mouse-adapted scrapie prion strain ^23-25^. Like RML prions^15^, ME7 prion rods are predominantly single-protofilament helical amyloid fibrils that coexist with paired protofilaments. Crucially, the fibrils of both mouse prion strains share the same underlying modular architecture, but with markedly altered topology. We identified conformationally conserved and variable regions in the N-terminal lobe, and a structurally congruent, but differently oriented, disulphide-stapled hairpin in the C-terminal lobe. ME7 and RML strain diversity appears to be linked to the divergent fold of the conformationally variable region that determines the orientations of the N- and C-terminal lobes, resulting in distinct helical assemblies.

## Results

### ME7 fibril morphologies resemble those of RML

ME7 prion rods were purified to ∼99% purity with respect to total protein from the brain of terminally infected C57Bl6 mice (Supplementary Fig. 1). We employed the same established method^23^ as for the analogous RML preparations^15^, which includes proteinase K (PK) treatment of crude brain homogenate and addition of phosphotungstate (PTA) polyanions that were shown to decorate RML prion fibrils^15^ without affecting the protofilament structure^15,16^. Mass spectrometry (MS) analyses showed that PK N-terminally truncates PrP monomers in ME7 rods at the same site as in RML rods^15^, leaving PrP subunits predominantly starting at residue 89 and extending to the C-terminus with intact GPI anchor. Prion infectivity of purified ME7 prions was measured using the Scrapie Cell End Point Assay^23,26,27^ (Supplementary Table 1) with titres consistent with previous findings^10,11,23^. As before^11,15^, prion rods were the only visible polymers in micrographs with the exception of occasional collagen fibres, amorphous aggregates or vesicles.

Among predominantly single protofilament fibrils (∼10 nm apparent diameter), distinctive paired assemblies with the apparent diameter of ∼20 nm were also observed in approx. 15% of the micrographs (Fig. 1a). As with the previously reported RML pairs, it remains unclear if or how PTA may influence the protofilament pairing^15^. Thus, here we focus on single ME7 protofilaments and on how they relate to the previously reported RML protofilaments^15^, whose structure is demonstrably not perturbed by PTA^16^.

**Figure 1.**
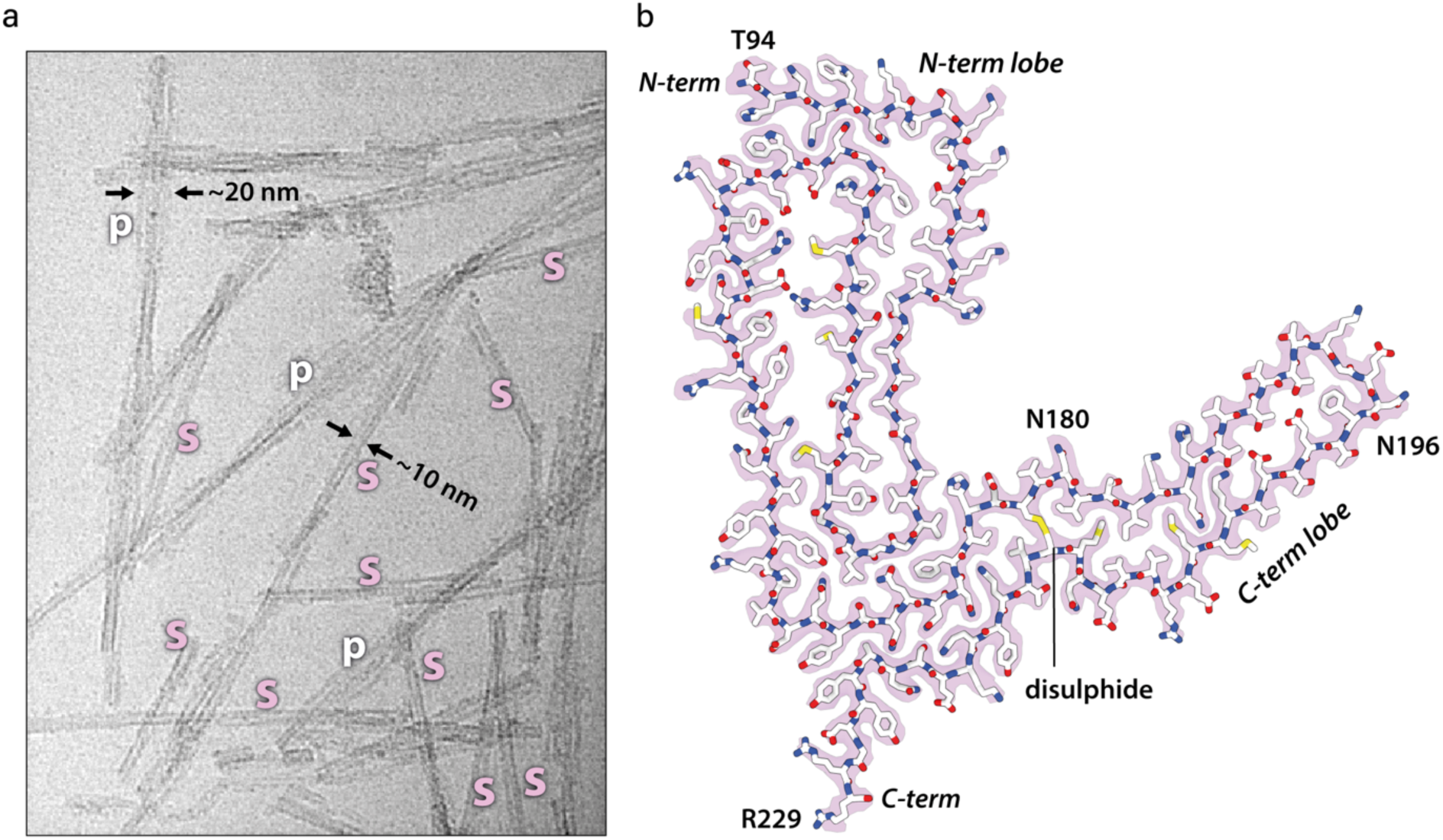
ME7 fibril morphologies and the atomic structure of their constituent PrP subunit. **a** Representative cryo-EM image (300 kV FEI Krios G3i, K3 camera) showing examples of single ME7 protofilaments (s) alongside their paired assemblies (p) with their approximate diameters. **b** Protein-only density of a single amyloid rung (pink) with the fitted atomic model of the mouse PrP chain shown with sticks coloured by heteroatom: C, white; N, blue; O, red; S, yellow. The start (T94) and end (R229) residues of the fitted polypeptide and both N-glycosylation sites are indicated.

We determined a 2.6 Å-resolution structure of the single ME7 protofilament and found that the fold of its constituent PrP subunit closely resembles that of the RML protofilament, including the double-hairpin N-terminal lobe and a single-hairpin disulphide-stapled C-terminal lobe, with four additional C-terminal residues stabilised as part of the amyloid core in the ME7 fibril (94-229) compared to the RML fibril (94-225) (Fig. 1b, Table 1 and Supplementary Fig. 2-3). We thus appended the remaining D226-R229 residues to the previously built model^15^ and then fitted and refined that C-terminally extended model in the cryo-EM density of ME7 (Fig. 1b and Table 1).

**Table 1.** Cryo-EM data collection, refinement and validation statistics.

|  | ME7<br>(EMDB-xxxx)<br>(PDB xxxx) |
| --- | --- |
| **Data collection and processing** |  |
| Magnification | 105,000 x |
| Voltage (kV) | 300 |
| Electron exposure (e<sup>-</sup>/Å<sup>2</sup>) | 49.4 |
| Defocus range (μm) | from -2.4 to -0.9 |
| Pixel size (Å) | 0.828 |
| Symmetry imposed | C1 |
| Initial particle images (no.) | 223,065 |
| Final particle images (no.) | 40,239 |
| Map resolution (Å) | 2.6 |
| FSC threshold | 0.143 |
| Map resolution range (Å) | 2.6-3 |
| **Refinement** |  |
| Initial model used (PDB code) | 7QIG |
| Model resolution (Å) | 2.6 |
| FSC threshold | 0.143 |
| Model resolution range (Å) | 121.7 - 2.6 |
| Map sharpening *B* factor (Å<sup>2</sup>) | -26.75 |
| **Model composition** |  |
| Non-hydrogen atoms | 6456 |
| Protein residues | 408 |
| Ligands | none |
| **B factors (Å<sup>2</sup>)** |  |
| Protein | 24.18-67.46 |
| **r.m.s. deviations** |  |
| Bond lengths (Å) | 0.002 |
| Bond angles (°) | 0.580 |
| **Validation** |  |
| MolProbity score | 1.49 |
| Clashscore | 2.79 |
| Poor rotamers (%) | 0 |
| **Ramachandran plot** |  |
| Favored (%) | 93.53 |
| Allowed (%) | 6.47 |
| Disallowed (%) | 0 |

### General comparison of ME7 and RML protofibrils

Akin to the previously reported RML protofilaments^15,16^, the ME7 protofilaments comprise a helical PIRIBS ribbon, where each two-lobed rib or rung is formed by a single PrP chain (Fig. 2a and b, Supplementary Fig. 3a and b). The left-handed twist of the ME7 fibril is slightly slower than that of the RML fibril (approx. -0.54° vs. -0.64° per rung, respectively), which results in a longer crossover distance (approx. 1585 vs. 1344 Å, respectively) (Fig. 2a). Spacing between the ME7 rungs was estimated at approx. 4.79 Å, while in RML at approx. 4.82 Å^15^ but given the uncertainty of the pixel size calibration for each magnification (different in each dataset) these spacings can likely be considered essentially the same (∼4.8 Å).

**Figure 2.**
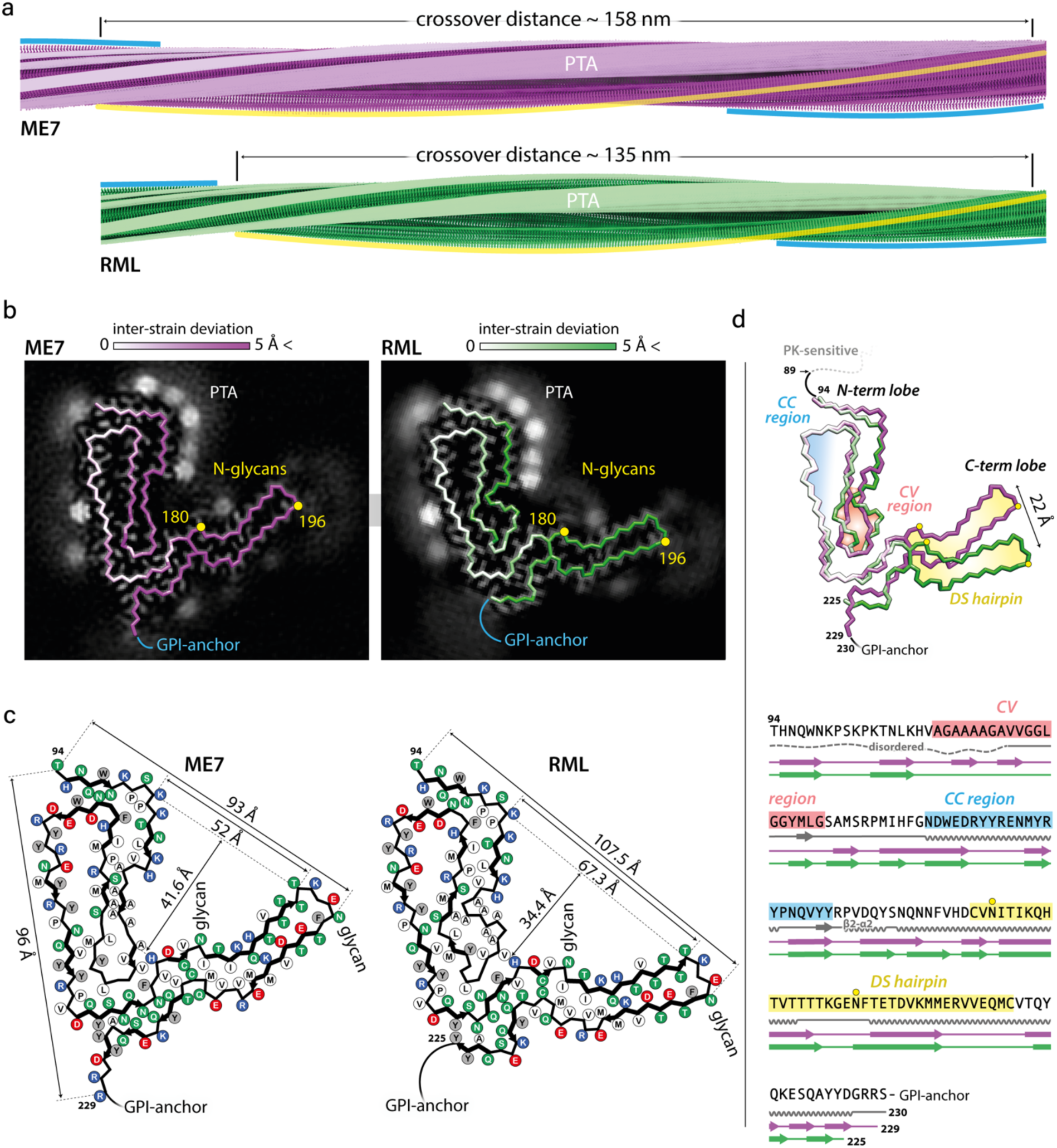
Comparison of ME7 and RML protofibrils and assignment of folding sub-domains in their core. **a** Rendered density views of the helix crossover or half-pitch (180° helical turn) distance, with annotated locations of phosphotungstate (PTA) polyanions (semi-transparent white), N196 (yellow) and GPI-anchor (blue). **b** Density cross-sections with overlaid PrP backbone models coloured by relative deviation based on global 3D alignment (UCSF Chimera; panel (d)), with annotations corresponding to panel (a). C-α atoms of both N-glycosylation sites (N180 and N196) are marked with yellow circles. **c** Diagrams of the PrP subunits with approximate dimensions of the inter-lobe grooves and the longest C-α distances in each model, measured between the indicated C-α atoms (dotted lines). Positions of amino acid side chains are shown with circles (positively charged, blue; negatively charged, red; neutral, green; hydrophobic, white; aromatic, grey) on either side of the backbone (black line). β-strands are indicated with thick black arrow-headed lines. **d** Top, superposition of PrP backbones coloured as in (b) and shaded according to the folding sub-domain assignment: CC, conformationally conserved; CV, conformationally variable; DS, disulphide-stapled. N-glycosylation sites are marked with yellow circles. PK-resistant core starts with residue 89. Bottom, mouse PrP sequence from the start of the amyloid core (T94) to the C-terminus (S230-GPI-anchor), with colour-coded folding sub-domain assignment. Secondary structure annotation for PrP^C^ (grey), ME7 fibril (magenta) and RML fibril (green) is included below the sequence (α-helix, zig-zag; β-sheet, arrow; disordered, undulated dashed line). Start and end residues are numbered and the β2-α2 region of PrP^C^ is indicated. N-glycosylation sites (N180 and N196) are marked with yellow circles.

The extra (non-protein) densities surrounding the N-terminal lobe of the ME7 reconstruction are consistent with PTA cages ([PW11O39]^7-^ at pH 7.8), previously seen in corresponding locations (basic residues) around RML rods purified with the same method^15^ (Fig. 2a-c, Supplementary Fig. 4a-b). The fainter extra densities around the C-terminal lobe of the ME7 reconstruction are also consistent with those previously seen around RML protofilaments^15^ and correspond to N180- and N196-linked glycans of variable occupancy and the flexible GPI-anchor at the C-terminus (Fig. 2b, Supplementary Fig. 4a). There are also likely at least two additional weak PTA-binding sites at residues K184 and K220 in the C-terminal lobe of ME7 and one at residue K184 in the C-terminal lobe of RML; both strains also show weak areas of unassigned density around the hydrophobic patch V202-M204 (compare Fig. 2b and c).

In each strain, the N180-glycan stems from the base of the C-terminal lobe and occupies the grove between the two lobes, which is significantly narrower and deeper in the ME7 fibril than that in the RML fibril (approx. 52 × 41.6 vs. 67.3 × 34.4 Å, respectively) (Fig. 2b-d, Supplementary Fig. 4a-b). The N196-glycan and GPI-anchor project outward in a roughly similar fashion in each fibril (Fig. 2a-c). Considering protein backbone, the widest dimension of the ME7 rung is shorter than that of RML (96 Å vs. 107.5 Å, respectively) and falls between the first (T94) and the last (R229) residue of the amyloid core (one residue away from the GPI-anchor), whilst that of RML falls between the first residue of the amyloid core (also T94) and N196 (the second glycosylation site) (Fig. 2c).

### Assignment of common PrP folding sub-domains in ME7 and RML protofibrils

Global 3D alignment of the PrP monomer from the ME7 fibril with that from the RML fibril reveals regions of conserved and of variable conformation (Fig. 2b and d). The N-terminal lobe can be sub-divided into two folding sub-domains on that basis. The tip of the first hairpin contains the Ala/Gly-rich sequence (A112-G130), which adopts a strikingly different conformation in the two strains (Fig. 2b-d, Supplementary Fig. 4c) and is hereafter designated the conformationally variable (CV) region. Crucially, this CV region interfaces with the C-terminal lobe, and thus, its conformation can impact the spatial arrangement of the two lobes (Fig. 2d). Conversely, the tip and the external half of the second hairpin (N142-Y162) have closely superimposable conformations in both strains (< 1 Å root mean square deviation, RMSD) (Fig. 2b and d and Supplementary Fig. 4c), we therefore designate this the conformationally conserved (CC) region. The disulphide-stapled (DS) hairpin of the C-terminal lobe, which harbours the two glycosylation sites is also nearly superimposable between the two strains (< 1 Å RMSD), although it is markedly displaced (by 22 Å at the tip) in the ME7 structure compared to the RML structure, due to the altered configuration of the CV region (Fig. 2d and Supplementary Fig. 4d).

### Other common structural regions in ME7 and RML protofibrils

K100-H110 region forms a major basic patch in the N-terminal lobe of both fibrils, and faces the N180-glycan in the groove of the fibril (Fig. 2b-c and 3a). This basic patch appears to be firmly stabilised by the first of the five major intra-chain hydrophobic clusters (mainly P101, P104, L108, V111, P136, I138 and F140), in both ME7 and RML fibrils (Fig. 2b, 3a and Supplementary Fig. 5). The second and third major hydrophobic clusters (mainly V121, Y126, L128, and L123, M127, V160, Y162, P164, respectively) appear to stabilise variable configurations of the CV region within the N-terminal lobe, whereas the fourth major hydrophobic cluster provides the third hydrophobic anchor point for the CV region at the inter-lobe interface in both protofibrils (Fig. 2c, 3a and Supplementary Fig. 5). This hydrophobic interface involves the N-terminal lobe’s residue V120 and the C-terminal lobe’s residues F174 and H176 in both RML and ME7 folds, and additionally N-terminal lobe’s residue A119 in the ME7 fold only (Fig. 2c and Supplementary Fig. 5). Finally, the fifth major hydrophobic cluster (I181, I183, M205, V208, V209, M212) precedes and encompasses the base of the DS hairpin (spanning both sides of the disulphide bond), likely conferring rigidity to the DS hairpin fold in both structures (Fig. 2c, 3a and Supplementary Fig. 5). Of note, the interaction of V175 with V214 in the ME7 protofibril is replaced by a less favourable interaction with T215 in RML (Fig. 2c).

### Distinct PIRIBS arrangement in ME7 and RML protofibrils

There are 15 inter-chain β-sheets (or PIRIB-sheets) in ME7 and RML protofibril structures, but their arrangement is not identical (Fig. 2c-d and 3b). The largest variation in the β-sheet distribution is seen in the CV region and the highest conservation of that is seen in the CC region (β-strands 6-8) and the DS hairpin (β-strands 10-12 in ME7 and 11-13 in RML) (Fig. 2c-d and 3b). Intervening regions also show variations in the PIRIBS arrangement (Fig. 2c-d and 3b), but the first two β-strands that accompany the major basic patch are conserved (Fig. 2c, 3a-b and Supplementary Fig. 5). The PIRIBS architecture defines the main longitudinal polar inter-chain interactions, the inter-rung hydrogen bonds, but besides the hydrophobic interactions described earlier, there are also lateral intra- and inter-chain hydrogen bonds that stabilise both folds (Supplementary Fig. 5).

Notably, all types of lateral associations between neighbouring strands are not exactly co-planar, but staggered by ∼half-rung distance along the helical axis (i.e. running between rungs). The most pronounced stagger (by nearly 1 rung) is seen at the inter-lobe interface of the RML protofibril (i.e. the N-terminal lobe of the rung i interacts with the C-terminal lobe of mainly the rung i + 1). In the ME7 protofibril this interface is more continuous, showing the standard ∼half-rung stagger (i.e. the N-terminal lobe of the rung i interacts equally with C-terminal lobes of the rungs i and i + 1) (Fig. 3b).

**Figure 3.**
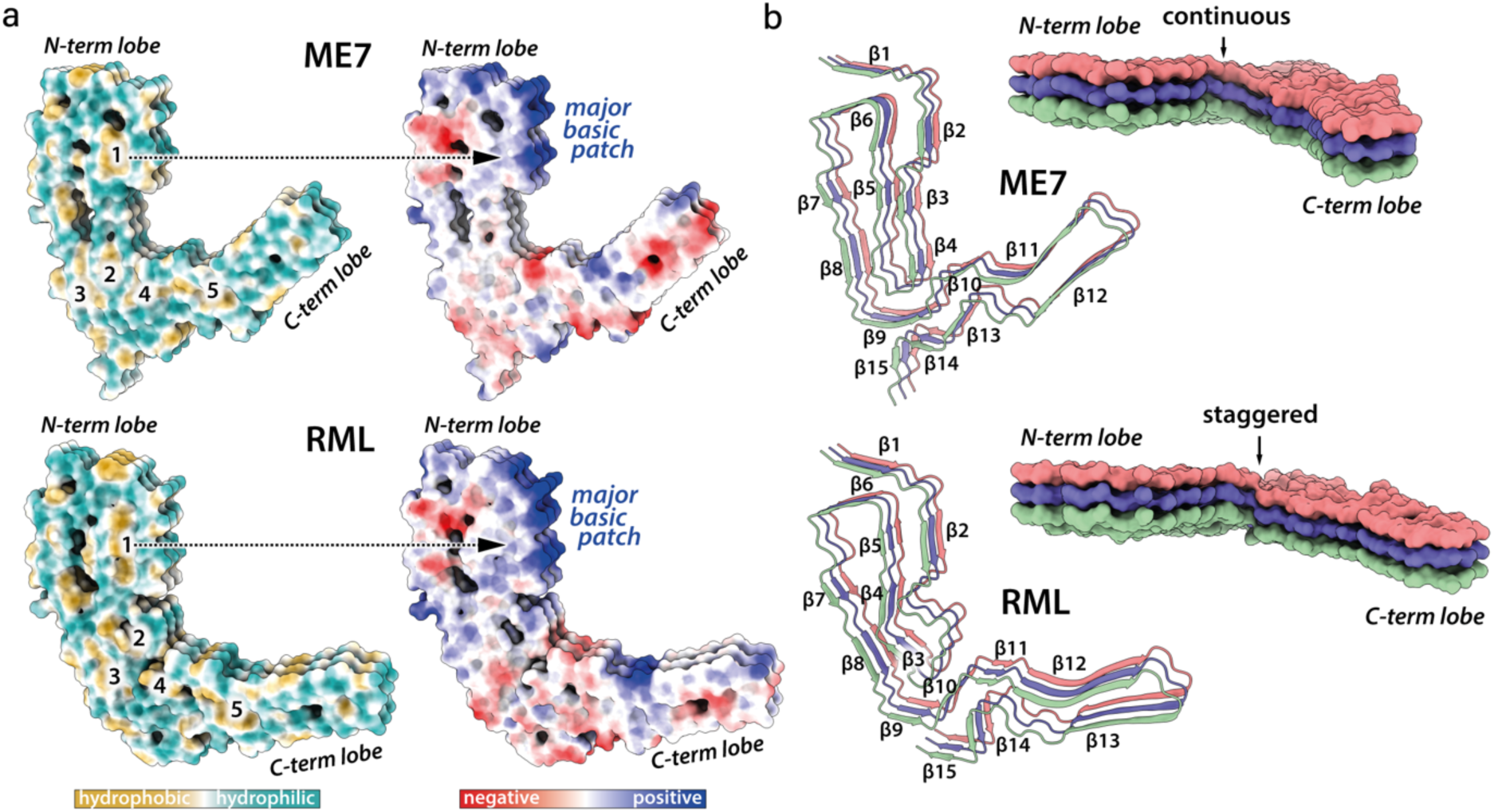
Hydrophobic and polar domains of mouse prion protofibrils and details of their PIRIBS arrangements. Models of three consecutive PrP rungs shown as: **a** solvent excluded surface coloured by hydrophobicity and by electrostatic charge distribution (ChimeraX), hydrophobic clusters are labelled 1-5; **b** ribbons with secondary structures and solvent-excluded surface models coloured by chain.

### N180 is more glycosylated than N196 in both ME7 and RML strains

By western blotting the ME7 strain has a greater proportion of di-glycosylated PrP than RML, whereas RML contains relatively more mono- and non-glycosylated PrP chains^23^ (Supplementary Fig. 1). However, both mono-glycosylated PrP variants appear as one band on a blot or a gel, and the relative abundance of the mono-180 vs. mono-196 PrP glycoform is not known in either strain. The N180-glycan faces the major basic patch in both protofibril structures, but it is closer to that patch in the ME7 protofibril (Fig. 2b-d). As the glycan chains are often sialylated, and more so in the ME7 than in the RML strain^28^, this raises the possibility of an electrostatic interaction between negatively charged sialylated N180-glycans and the positively charged residues of the basic patch that may promote mono-180 glycoform incorporation. To inform on the proportions of mono-180 and mono-196 glycoforms in the ME7 and RML protofibrils we used mass spectrometry (MS) to quantify the relative abundance of all N180- and N196-glycans (Fig. 4). Purified rods were first denatured and then treated with PNGase F to enzymatically remove all N-linked glycans, a process which results in the conversion of the underlying asparagine residue to an aspartic acid^29^. This resultant N-to-D ‘quasi-mutation’ at each glycosylation site acts as a surrogate marker for prior glycosylation that can be readily quantified by MS. Complete de-glycosylation of PrP from purified *ex vivo* prion rod preparations was confirmed by western blot, wherein total PrP was observed as a single band (Supplementary Fig. 6a). In-gel trypsinisation of this band produced the following peptides spanning the N180- and N196-glycosylation sites: ^156^YPNQVYYRPVDQYSNQNNFVHDCV-(N/D)-ITIK^184^ and ^194^GE-(N/D)-FTETDVK^203^ and the relative abundance of the D-isoform of each peptide was taken as a representation of total glycan occupancy at that site. (Fig. 4a and Supplementary Fig. 6b-c). We found that in both strains the proportion of N180- and N196-glycans are equally skewed towards the N180 site (ratio of ∼1.2), with the ME7 strain showing more overall glycosylation, as expected from its higher content of di-, and lower content of non-glycosylated PrP chains compared to RML (Fig. 4b). Thus, despite a shorter distance between the N180 glycosylation site and the major basic patch in the ME7 protofibril compared to the RML protofibril this does not appear to influence incorporation of mono-180 vs. mono-196 PrP glycoforms.

**Figure 4.**
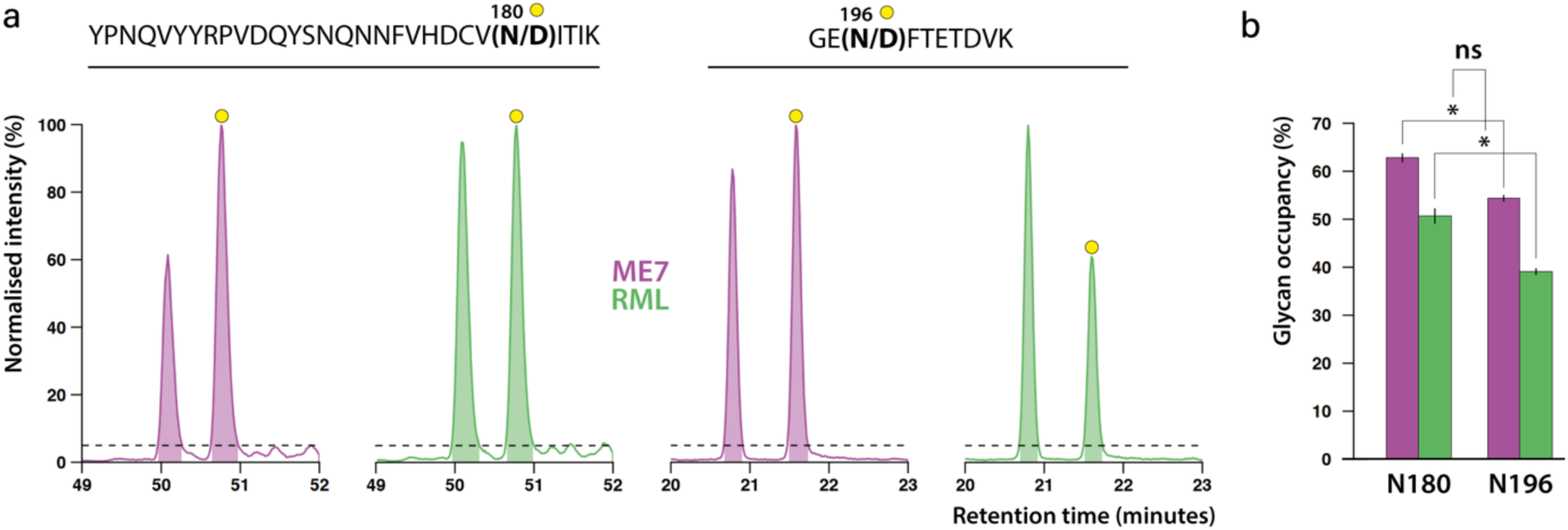
Glycan occupancy at N180 and N196 glycosylation sites. **a** Normalised LC-MS total ion current peaks representing relative abundances of non-glycosylated (unlabelled peaks) and de-glycosylated (yellow circles) sites. The N180 and N196 site are represented by PrP peptides shown, as produced by in-gel trypsinisation of denatured, PNGase-treated and electrophoresed purified ex vivo prion rods. Conversion of N-glycosylated residues to D by PNGase F increases the retention time of the resultant peptides on the LC column, enabling their separation and quantification, as indicated by shaded areas under the peaks (dashed line indicates the peak width cut-off point at 5% maximal signal intensity). **b** Bar graph summarising glycan occupancy at each glycosylation site, colour-coded as in (a). Each bar represents the average ratio of non-glycosylated and de-glycosylated sites from three independent experiments (error bar, standard deviation). *, statistically significant (T-test: p < 0.05); ns, not significant. See also Supplementary Fig. 6.

## Discussion

The infectious RML and ME7 mouse prion protofibril structures compared here show the same modular architecture, comprising two conformationally congruent modules (the CC region in the N-terminal lobe and the DS hairpin in the C-terminal lobe) connected by a conformationally variable module (the CV region) at the core of the assembly (Fig. 5). A similar overall architecture is seen in the hamster 263K protofibril^14-16^ (Supplementary Fig. 7). Comparison of the three folds suggests that the low-complexity (Gly/Ala-rich) region of PrP (residues 112-130) (Fig. 5b) that comprises the CV region is likely amenable to a greater range of misfolded configurations, whereas other more complex regions of sequence that comprise the CC region and DS hairpin (Fig. 5b), may be more conformationally restricted (Supplementary Fig. 7). Of note, in all three folds the inter-lobe interface involves interaction of residues of the CV region with residues that comprise the β2-α2 loop in the normal PrP^C^ fold (residues 165-175)^30-32^ (Fig. 2d and Supplementary Fig. 7). Variation of amino acid sequence within the β2-α2 loop has dramatic effects on numerous interspecies prion transmission barriers^33-37^ and the CV region contains the 127G/V and 129M/V polymorphisms of human PrP that profoundly affect prion disease susceptibility, phenotype and strain selection^2-4,21,38-43^. Thus the inter-lobe interface appears to be a critical structural determinant of conformational selection linking strains and prion transmission barriers. While structures of protofibrils from other prion strain/host combinations are clearly required to inform on the generalisability of these new findings, these data now suggest an initial structural framework underpinning prion strain diversity in mammals.

**Figure 5.**
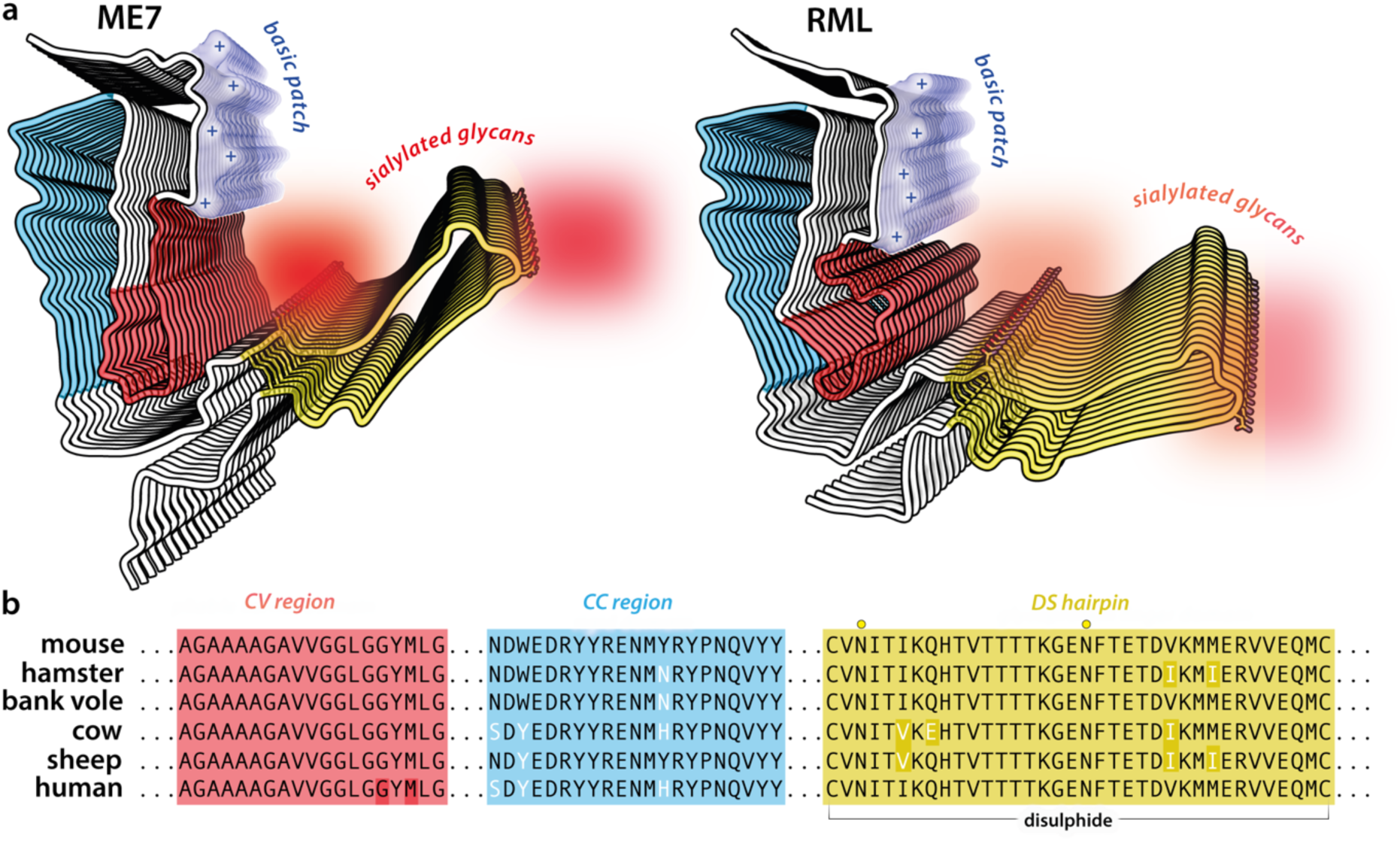
Summary of structural differences between ME7 and RML prion protofibril structures. **a** Licorice backbone models (ChimeraX) of ME7 and RML protofibrils coloured according to PrP^Sc^ folding sub-domains (blue, CC region; red, CV region; yellow, DS hairpin; white, intervening regions). Positive charge of the major basic patch (transparent surface) is indicated with + signs, and the sialoglycan occupancy corresponds to the intensity of red shading. The N-glycosylation sites (N180 and N196 side chains) and the disulphide bond are shown as sticks and coloured by heteroatom (C, yellow; N, blue; O, red; S, yellow). **b** Alignment of PrP sequences from different mammals focused on the PrP^Sc^ folding sub-domains. CV, conformationally variable; CC, conformationally conserved; DS, disulphide-stapled. Human residues 127 and 129 in the CV region (highlighted with dark red) are polymorphic (G/V and M/V, respectively).

Notably, the K100-H110 sequence is part of the disordered N-terminal domain in PrP^C 30- 32^(Fig. 2d), but in both mouse (ME7 and RML) protofibrils and in the hamster 263K protofibril that amino acid stretch forms a common physicochemical motif, the major basic patch, around the second β-strand of the N-terminal lobe (compare Figs. 2c, 3a-b, 5a and Supplementary Fig. 7) ^14-16^. This basic patch faces the N180-glycan in both RML and ME7 mouse protofibrils and also in the hamster protofibril^14-16^ and may therefore interact with sialylated N180-glycans. However, PTA polyanions used in our *ex vivo* prion fibril preparations clearly line the major basic patch of both mouse prion protofibrils (Fig. 2b), and in turn change its charge from positive to negative. This change does not unify the geometry of ME7 and RML protofibrils, suggesting that the strain-specific topology of the N- and C-terminal lobes in mature prion strains is locked by the PIRIBS architecture alone and does not depend on long-range sialylated glycan-protein interaction. However, as we recently proposed, recruitment of particular PrP isoforms into nascent prion fibrils at the inception of a prion strain might critically determine the configuration of the fold^15^.

Our MS analyses showed that variable distance between the major basic patch and the N180-glycosylation site does not impact the ratio of mono-glycosylated PrP variants in ME7 and RML protofibrils. Moreover, stacking of solely di-glycosylated PrP chains in the PIRIBS architecture does not appear energetically prohibitive^44^. It therefore remains unclear how the single protofibril structures alone might determine particular PrP glycoform compositions. We propose that the paired fibril assemblies observed in our *ex vivo* preparations may have a critical role in determining strain-specific PrP glycoform signatures^10,11,15^.

These pairs currently represent a minor sub-population (10-15%) of assemblies observed in our RML and ME7 fibril preparations. However, we do not know how different prion purification methods may distort the true (*in vivo*) content of the pairs. For example, PTA cages are found adjacent to various pairing interfaces and may be disruptive^15^. Whether relatively harsh purification conditions used by others, for example 1.7 M NaCl^14^, may also be disruptive remains to be established. Structural investigation of the pairs purified without PTA is currently ongoing and may shed more light on how prion strain-specific glycoform ratios are generated.

## Methods

### Research governance

Prion purification, cell-based prion bioassay and preparation of cryo-EM grids was conducted at UCL in microbiological containment level 3 or level 2 facilities with strict adherence to safety protocols. Work with infectious prion samples at Birkbeck College London was performed using dedicated sample holders and equipment with strict adherence to safety procedures and local risk assessment. Prion samples were transported between laboratories in packaging conforming to UN 3373 Biological Substance, Category B specifications. Frozen brains from mice with clinical prion disease were used to generate purified prion samples. These brain samples were generated by us as part of a previous study^23^ in which work with animals was performed in accordance with licences approved and granted by the UK Home Office (Project Licences 70/6454 and 70/7274) and conformed to University College London institutional and ARRIVE guidelines. All experimental protocols were approved by the Local Research Ethics Committee of UCL Queen Square Institute of Neurology/National Hospital for Neurology and Neurosurgery.

### Preparation of purified ME7 prion rods

Prion-infected brain homogenate was prepared by homogenizing 30 brains from female C57Bl/6 mice terminally-infected with the ME7 prion strain in Dulbecco’s phosphate buffered saline lacking Ca^2+^ or Mg^2+^ ions (D-PBS; Gibco) to produce a pool of 130 ml 10% (w/v) ME7 brain homogenate (designated I21487) using methods described previously^23^. Purification of ME7 prion rods was performed as described previously^23^ with the exception first implemented in Manka et al. (2022)^15^ that initial protease digestion was performed using PK in the place of pronase E. 200 μl aliquots of 10% (w/v) ME7 brain homogenate were dispensed into standard 1.5 ml microfuge tubes with screw cap and rubber O ring. Typically, 12 tubes were processed at a time. Samples were treated with 2 μl of 5 mg/ml PK prepared in water (to give 50 μg/ml final protease in the sample) and incubated for 30 min at 37°C with constant agitation, after which digestion was terminated by addition of 4.1 μl of 100 mM 4-(2-Aminoethyl) benzenesulfonyl fluoride hydrochloride (AEBSF) to give 2 mM final concentration in the sample. 206 μl of 4% (w/v) sarkosyl (Calbiochem) in D-PBS and 0.83 μl of Benzonase (purity 1; 25,000 U/ml) were then added to give final concentrations in the sample of 2% (w/v) and 50 U/ml, respectively. Following incubation for 10 min at 37°C, 33.5 μl of 4% (w/v) sodium phosphotungstate (NaPTA) prepared in water pH 7.4 was added to give a final concentration of 0.3% (w/v) in the sample. After incubation for 30 min at 37°C the samples were adjusted (and thoroughly mixed) with 705.3 μl of 60% (w/v) iodixanol and 57.2 μl of 4% (w/v) NaPTA prepared in water pH 7.4, to give final concentrations in the sample of 35% (w/v) and 0.3% (w/v), respectively. After centrifugation for 90 minutes at 16,100*g* the sample separates into an insoluble pellet fraction (P1), a clarified supernatant (SN1) and a buoyant, partially flocculated, surface layer (SL). 1 ml of SN1 was carefully isolated from each tube taking extreme care to avoid cross contamination with either P1 or SL. SN1 was filtered using an Ultrafree-HV microcentrifuge filtration unit (0.45 μm pore size Durapore membrane, Millipore, Prod. No. UFC30HV00). This was accomplished by loading 500 μl aliquots of SN1 and centrifugation at 12,000*g* for 30 sec using one filtration unit per ml of SN1. 480 μl aliquots of filtered SN1 were transferred to new 1.5 ml microfuge tubes and thoroughly mixed with an equal volume of 2% (w/v) sarkosyl in D-PBS containing 0.3% (w/v) NaPTA pH 7.4 and incubated for 10 min at 37°C. Samples were then centrifuged for 90 min at 16,100*g* to generate an insoluble pellet fraction (P2) and a clarified supernatant (SN2). SN2 was carefully removed and discarded, after which each P2 pellet was resuspended in 10 μl of 5 mM sodium phosphate buffer pH 7.4 containing 0.3% (w/v) NaPTA and 0.1% (w/v) sarkosyl. In order to avoid unnecessary aggregation of the purified rods arising from repeated rounds of centrifugation the final two wash steps detailed in Wenborn et al. (2015)^23^ were replaced with a single wash. Resuspended P2 pellets were pooled and mixed with 1.0 ml of 5 mM sodium phosphate buffer pH 7.4 containing 0.3% (w/v) NaPTA and 0.1% (w/v) sarkosyl and samples centrifuged at 16,100*g* for 30 min to generate a clarified supernatant (SN3) and an insoluble pellet fraction (P3). SN3 was carefully removed and discarded and final P3 samples were typically resuspended to a concentration of 150-200X relative to the starting volume of 10% (w/v) brain homogenate from which they were derived, prior to loading on to EM grids (see below).

Prion infectivity of brain homogenates or purified samples was measured using the Scrapie Cell End Point Assay (SCEPA)^23,26,27^ using LD9 cells (an established cell line; that was a gift from Professor Charles Weissmann and originally derived from murine L929 fibroblasts supplied by the American Type Culture Collection^27^). Every experiment included concomitant assay of a serial dilution of RML prions of known prion titre determined from rodent bioassay. 10% (w/v) RML brain homogenate I6200 was used as the standard and reported a prion titre of 10^7.3 + 0.5^ (mean + s.d.) intracerebral LD50 units/ml when endpoint titrated six times in Tg20 mice that overexpress mouse PrP on a *Prnp*^o/o^ background^23^. PrP concentrations in purified samples were measured by ELISA as described previously^23^.

### SDS-PAGE, silver staining and western blotting

Samples were prepared for SDS-PAGE using NuPage 4X LDS buffer and 10X Reducing Agent (Thermo Fisher) according to the manufacturer’s instructions followed by immediate transfer to a 100°C heating block for 10 min. Electrophoresis was performed on NuPage 12% Bis-Tris protein gels (Thermo Fisher), run for 60 min at 200 V, prior to electroblotting to Immobilon P membrane (Millipore) for 16 h at 15 V. Membranes were blocked in 1X PBS (prepared from 10X concentrate; VWR International) containing 0.05% (v/v) Tween 20 (PBST) and 5% (w/v) non-fat dried skimmed milk powder and then probed with 0.2 μg/ml anti-PrP monoclonal antibody ICSM35 (D-Gen Ltd) in PBST for at least 1 h. After washing (1 h with PBST) the membranes were probed with a 1:10,000 dilution of alkaline-phosphatase-conjugated goat anti-mouse IgG secondary antibody (Sigma-Aldrich, Cat No A2179) in PBST. After washing (1 h with PBST and 2 × 5 min with 20 mM Tris pH 9.8 containing 1 mM MgCl2) blots were incubated for 5 min in chemiluminescent substrate (CDP-Star; Tropix Inc) and visualized on Biomax MR film (Carestream). SDS-PAGE gels (prepared as above) were silver stained using the Pierce Silver Stain Kit (Thermo Fisher) according to the manufacturer’s instructions. Gels were photographed on a light box using a Nikon Coolpix P6000 digital camera. Typical sample loadings for western blotting or silver staining correspond to purified material derived from 10 μl or 100 μl of 10% (w/v) RML prion-infected brain homogenate per lane, respectively.

### ME7 sample preparation for cryo-EM

ME7 prion rods purified from 2.4 ml 10% (w/v) ME7-infected brain homogenate were resuspended from the P3 pellet (see above) in 10-20 μl 5 mM sodium phosphate buffer Ph 7.4 containing 0.1% (w/v) sarkosyl and 4 μl of the suspension was applied directly to a glow-discharged C-flat™ Holey Carbon CF-2/2-4C Cu 400 mesh cryo-EM grid (Electron Microscopy Sciences) or Quantifoil R2/2 Cu 300 mesh grids in the chamber of the Leica GP2 plunging robot. The chamber was set to 20°C and 40% humidity. After 10 s incubation, the grids were blotted for 3 s (with an additional 2 mm push) and plunge-frozen in liquid ethane maintained at -183°C.

### Cryo-EM data collection

Cryo-micrographs were acquired at Birkbeck College London, on a 300 kV Krios G3i microscope (FEI/Thermo Fisher) with a post-GIF (20 eV slit) K3 detector (Gatan) operated in super-resolution bin 2x mode at 105,000 nominal magnification. The final (post-binning) magnified pixel size was 0.828 Å. The dose rate was ∼19.9 e-/Å^2^/s during 2.5-s exposures, resulting in a total dose of ∼49.75 e-/Å^2^ on the specimen. The exposures were collected automatically at 5 shots per grid hole, with fast acquisition (up to ∼370 images/hr), using the EPU 2 software (FEI/Thermo Fisher), at defocus ranging from 2.4 to 0.9 μm, and fractionated into 50 movie frames.

### Cryo-EM image processing and 3D reconstruction

All image processing except filament picking was done within the Relion 4.0-beta framework^45^. We used Relion’s implementation of the MotionCor2 algorithm to align movie frames. The contrast transfer function (CTF) parameters were estimated with CTFFIND4^46^. We then trained the deep learning package crYOLO^47,48^ to pick ME7 filaments using 100 example micrographs, as previously reported^15^. We imported the coordinates into Relion and extracted images of prion rod segments with different box sizes (ranging from 1024 to 384 pixels) to perform reference-free 2D classifications. Optimal 2D class averages and segments were selected for further processing and used to *de novo* generate an initial 3D reference with relion_helix_inimodel2d programme^49^, using an estimated rise of 4.79 Å and helical twist according to the observed crossover distances of the filaments in the 2D class averages. After 3D classification and 3D auto-refinement, we obtained a 3D reconstruction of the ME7 protofibril at 2.9 Å resolution. Subsequent Bayesian polishing^50^ and CTF refinement^51^ were performed to further improve the resolution of the reconstruction to 2.6 Å, according to the 0.143 FSC cut-off criterion (Supplementary Fig. 3).The final 3D map was sharpened with a generous, soft-edged solvent mask at 10% of the height of the box using the computed B-factor value of -26.75 Å^2^. The sharpened map was used for the subsequent atomic model building and refinement. The absolute hand of the helical twist was determined directly from the map through resolved densities of the carbonyl oxygen atoms of the polypeptide backbone^49^. The local resolution calculation was performed by LocRes in Relion 4.0-beta with solvent mask over the entire map.

### Atomic model building and refinement

A single subunit repeat was extracted from the cryo-EM map of the ME7 protofibril in UCSF Chimera^52^. A single PrP chain from the previously determined atomic model of the RML fibril structure (pdb id: 7QIG)^15^ was C-terminally extended by addition of D226-R229 residues and fitted to the extracted ME7 density in Coot^53^. The initially fitted atomic model was then copied and fitted into 3 consecutive subunits in the ME7 map and the map was zoned around the atomic coordinates in UCSF Chimera^52^. The 3-rung map and model were placed in a new unit cell with P1 space group for subsequent model refinement using default settings in phenix.real_space_refine^54^ and REFMAC5^55^. Model geometry was evaluated using MolProbity^56^ (http://molprobity.biochem.duke.edu/) after each round of refinement, and problematic or poorly fitting regions in the model were manually adjusted using Coot^53^ and Isolde^57^ (within ChimeraX^58^). This process was repeated until a satisfactory level of model:map agreement with acceptable model stereochemistry was achieved (Table 1).

### Structure analyses and presentation

Analyses and visualisations of the cryo-EM density map and the models compared in this study were done using UCSF Chimera^52^ and ChimeraX^58^.

### Determination of N-terminal PK-cleavage sites by mass spectrometry

N-terminal PK-cleavage sites in PrP were determined as described previously^15^. Briefly, purified RML or ME7 fibils were prepared as above and subjected to SDS-PAGE separation using NuPAGE 12% Bis-Tris mini protein gels (Thermo Fisher) according to the manufacturer’s instructions. Gel sections spanning all three PrP glycoforms were excised, reduced with 100 μM tris(2-carboxyethyl)phosphine and alkylated with 200 μm iodoacetamide prior to N-terminal labelling with 6 mM N-Succinimidyloxycarbonylmethyl tris(2,4,6-trimethoxyphenyl)phosphonium bromide, (TMPP-Ac-OSu) (Sigma) for 1 h at 22°C in 100 mM HEPES buffer pH 8.2. Gel-pieces were then washed thoroughly and subjected to overnight trypsin digestion at a working concentration of 2.5 μg/ml. The following day, tryptic digest peptides were recovered by gel extraction, as described by Shevchenko et al. (2006)^59^. Peptide analysis was performed by liquid chromatography mass-spectrometry, using an Acquity I-Class UPLC system coupled to a Xevo G2-XS Q-ToF mass spectrometer (Waters). Data was collected in MSe acquisition mode using concurrent low- and high-collision energy functions with 5V and 15-45V of collision energy respectively. Peptide sequence assignments were made using ProteinLynx Global Server 3.0.3 (Waters) against a species-specific reference proteome (Uniprot UP000000589, mus musculus) and optionally allowing for N-terminal amino-group derivatization by TMPP (+572.1811 Da). Extracted ion chromatograms were generated for each TMPP-labelled peptide and relative abundance was determined from their respective peak areas.

### Determination of glycosylation site occupancy by mass spectrometry

Glycosylation site occupancy in purified *ex vivo* prion preparations was determined by enzymatic de-glycosylation of PrP using PNGase F and subsequent analysis by mass spectrometry to monitor for the conversion of asparagine to aspartic acid that occurs during glycan hydrolysis. Briefly, purified RML or ME7 fibrils were prepared as above and digested using recombinant PNGase F (New England Biolabs) according to the manufacturer’s instructions. Samples were then subjected to SDS-PAGE separation using NuPage 12% Bis-Tris mini protein gels (Thermo Fisher) according to the manufacturer’s instructions and gel sections comprising the de-glycosylated band of PrP were excised. Tryptic digest peptides were prepared as described by Shevchenko et al. (2006)^59^ prior to analysis by liquid chromatography-mass spectrometry, using an Acquity I-Class UPLC system coupled to a Xevo G2-XS Q-ToF mass spectrometer (Waters). Survey data was collected in MSe acquisition mode in order to assess the distribution of ionisation states of the relevant non-glycosylated and de-glycosylated peptides, prior to MSMS analysis of the predominant ion species. Peptide identities were confirmed by MSMS parent ion and fragmentation spectrum peak assignments. Quantification was performed by determining chromatographic peak area of MSMS parent ion traces using a threshold intensity of 5%. Glycosylation site occupancy was expressed as the peak area of the de-glycosylated peptide as a proportion of the summed peak area of both de-glycosylated and non-glycosylated peptides.

## Acknowledgements

This work was funded by the core award to the MRC Prion Unit at UCL from the UK Medical Research Council (MC_U12316055 and MC_UU_00024/5). EM data collection was supported by grants from the Wellcome Trust (202679/Z/16/Z, 206166/Z/17/Z, 106249/Z/14/Z). We are very grateful to Dr Natasha Lukoyanova and Dr Shu Chen at Birkbeck College for EM support and Damian Johnson and Peter King for infrastructure support at UCL.

## Supplementary Information

**Supplementary Table 1.** Determination of prion infectivity titre in cell culture.

| Dilution | Positive Wells (w/w<sub>0</sub>) |  |  | Percentage Positivity |  |  |
| --- | --- | --- | --- | --- | --- | --- |
|  | RML BH<br>I6200* | ME7 BH<br>I21487 | Purified<br>ME7 (1X)† | RML BH<br>I6200* | ME7 BH<br>I21487 | Purified<br>ME7 (1X)† |
| 1.10<sup>-5</sup> | 24 / 24 | 24 / 24 | 24 / 24 | 100.0% | 100.0% | 100.0% |
| 3.10<sup>-6</sup> | 24 / 24 | 24 / 24 | 24 / 24 | 100.0% | 100.0% | 100.0% |
| 1.10<sup>-6</sup> | 20 / 24 | 24 / 24 | 15 / 24 | 83.3% | 100.0% | 62.5% |
| 3.10<sup>-7</sup> | 5 / 24 | 16 / 24 | 5 / 24 | 20.8% | 66.7% | 20.8% |
| 1.10<sup>-7</sup> | 2 / 24 | 9 / 24 | 1 / 24 | 8.3% | 37.5% | 4.2% |
| 3.10<sup>-8</sup> | 0 / 24 | 2 / 24 | 0 / 24 | 0.0% | 8.3% | 0.0% |
| 1.10<sup>-8</sup> | 0 / 24 | 1 / 24 | 0 / 24 | 0.0% | 4.2% | 0.0% |
| NI | 0 / 24 | 0 / 24 | 0 / 25 | 0.0% | 0.0% | 0.0% |
| log TCIU ml<sup>-1</sup>‡ |  |  |  | 5.41 | 6.09 | 5.51 |

(w/w<sub>0</sub>), the number of prion infected wells (w) versus the total number of inoculated wells (w<sub>0</sub>); BH, brain homogenate; TCIU, tissue culture infectious units; NI, non-prion inoculated
\* Prion titre of 10<sup>7.3 ± 0.5</sup> (mean + s.d., n=6) intracerebral LD<sub>50</sub> units/ml in Tg20 mice<sup>1</sup>
† Purified material was resuspended to the same volume as the 10% (w/v) BH from which it was derived
‡ TCIU calculated according to the Poisson distribution<sup>2</sup>

### Supplementary Figures

**Supplementary Figure 1.**
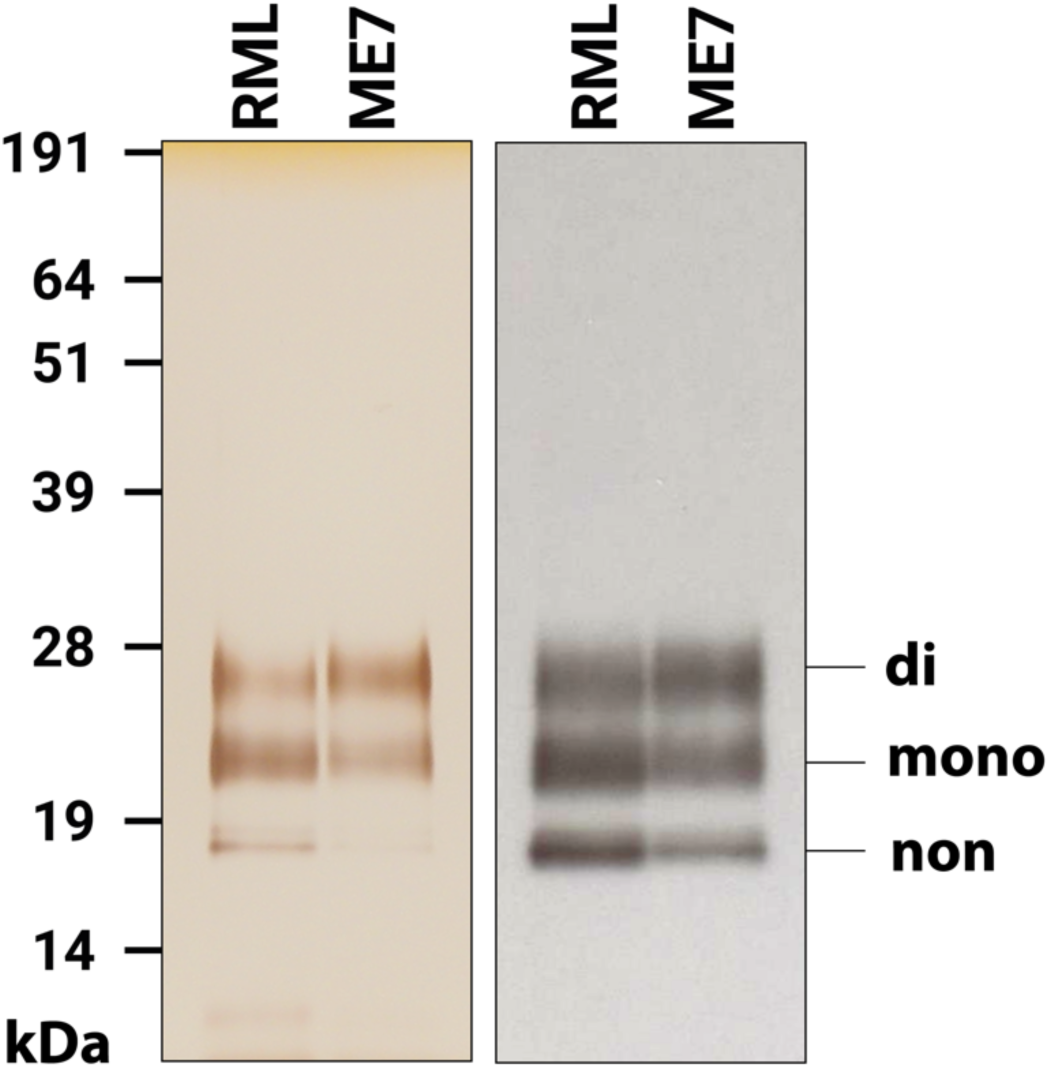
Silver-stained SDS-PAGE (left) and western blot (right) of purified ME7 and RML rods. Labelling shows the migration positions of di-, mono- and non-glycosylated PrP. The samples were prepared as described in Methods.

**Supplementary Figure 2.**
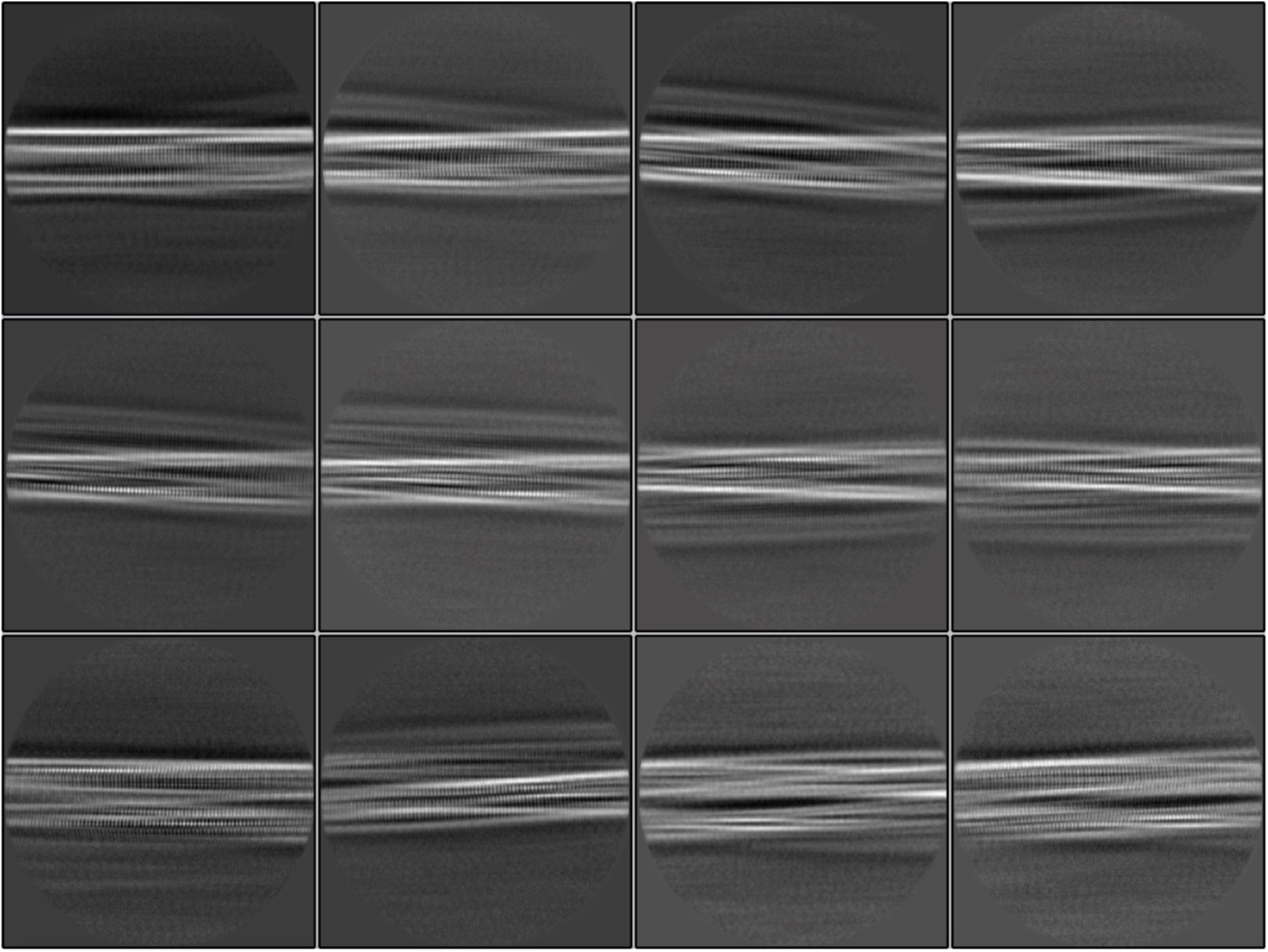
2D classification of ME7 prion fibrils. Gallery of representative 2D classes (box size: 397.5 × 397.5 Å).

**Supplementary Figure 3.**
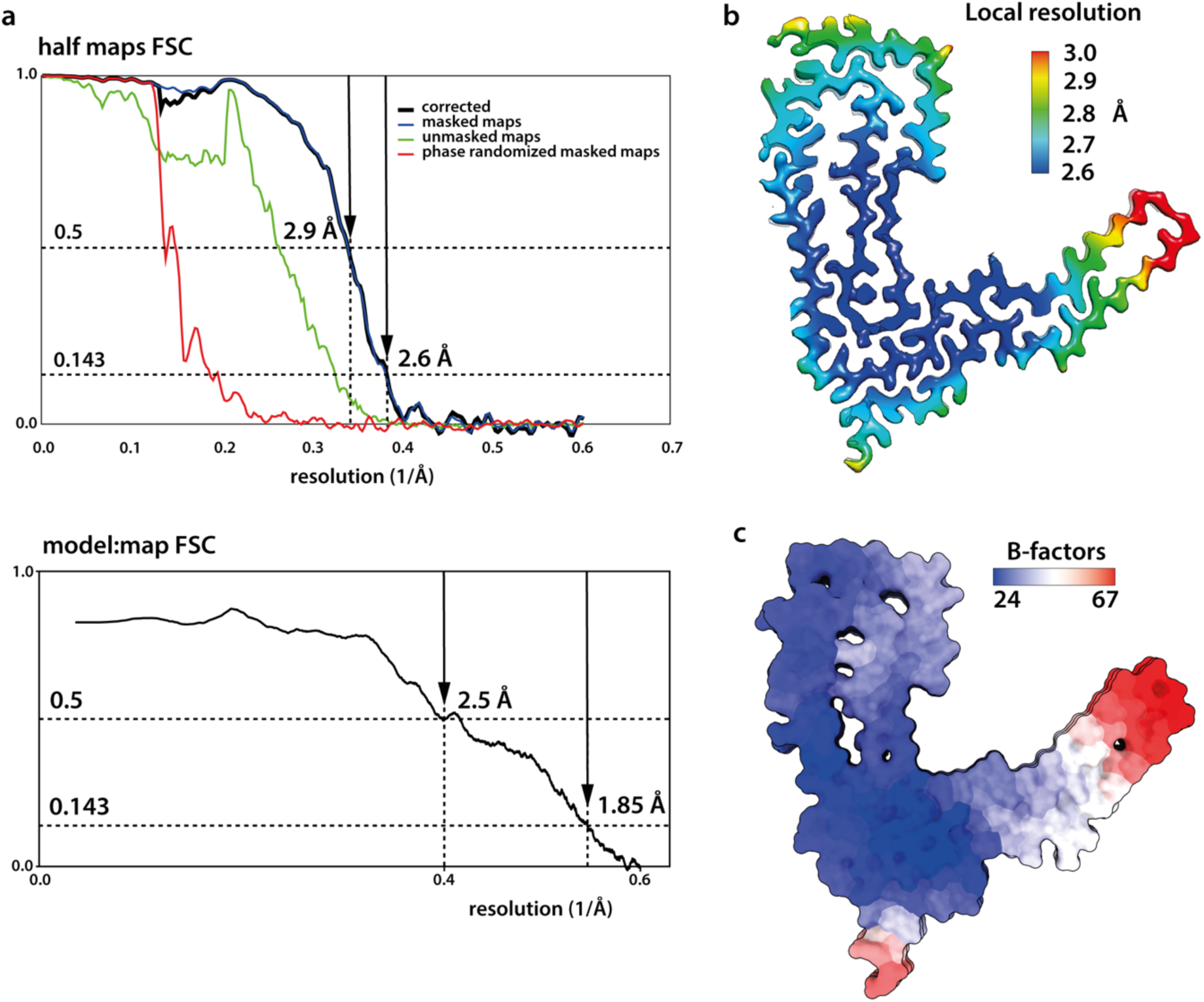
Quality of the ME7 map and model. **a** Top, Fourier Shell Correlation (FSC) plots for independent half-reconstructions, determined in Relion 4.0-beta. The final plot (black line) is corrected for overfitting with high-resolution noise substitution. Bottom, FSC plot for the model:map fit. **b** Local resolution of the protein density calculated with Relion 4.0-beta LocRes. **c** Atomic B-factor values colour-coded on the solvent-excluded model surface.

**Supplementary Figure 4.**
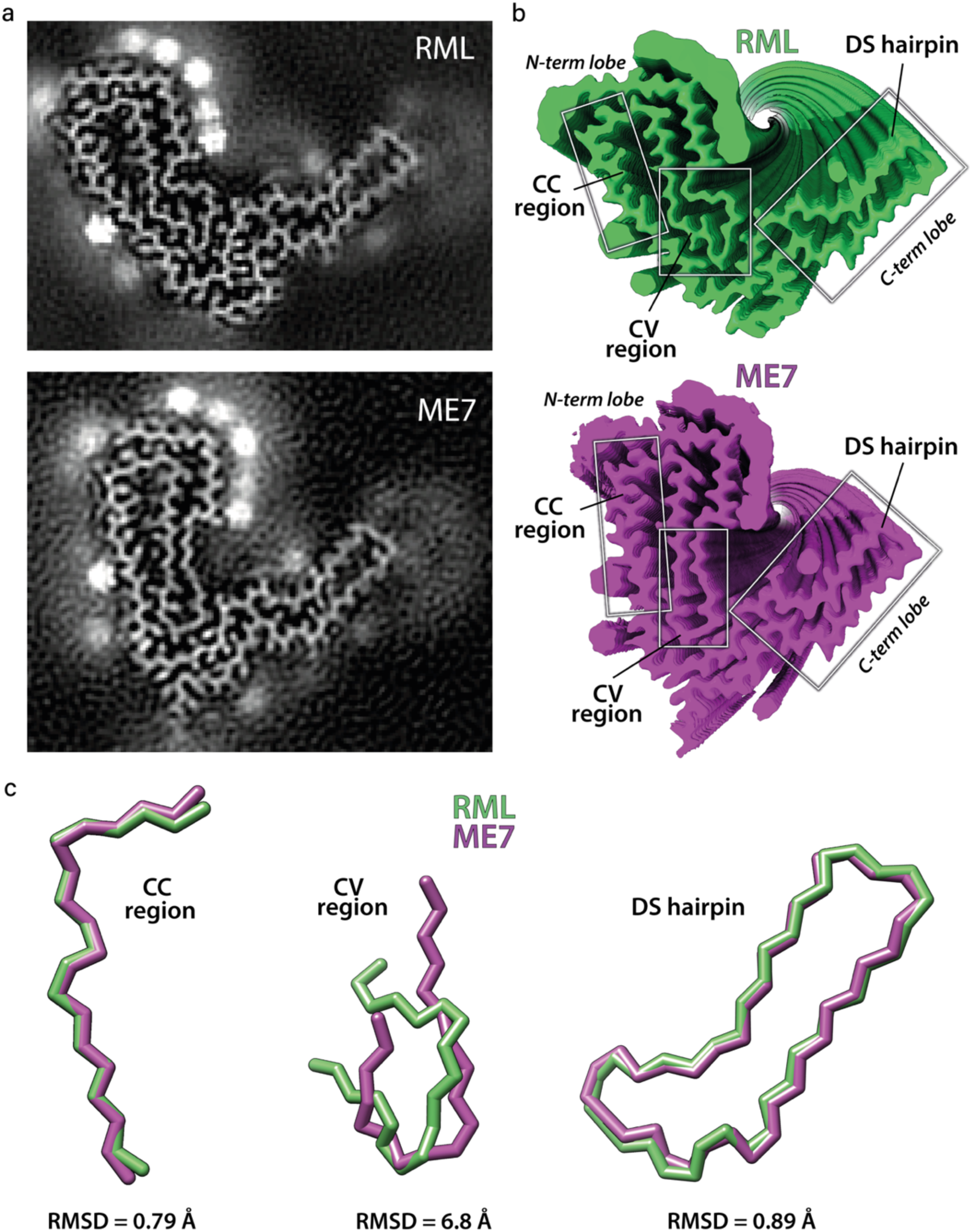
Comparisons between cryo-EM maps and atomic models of mouse prion fibrils. **a** Density cross-sections (pixel size: RML, 1.067 Å; ME7, 0.828 Å) showing the protein core and the non-protein extra densities. **b** Top views of rendered reconstructions with indicated common folding sub-domains. **c** backbone alignments of the common folding sub-domains and the root mean square deviations (RMSD) between all of their respective atom pairs, as calculated using UCSF Chimera^3^. CC, conformationally conserved; CV, conformationally variable; DS, disulphide-stapled.

**Supplementary Figure 5.**
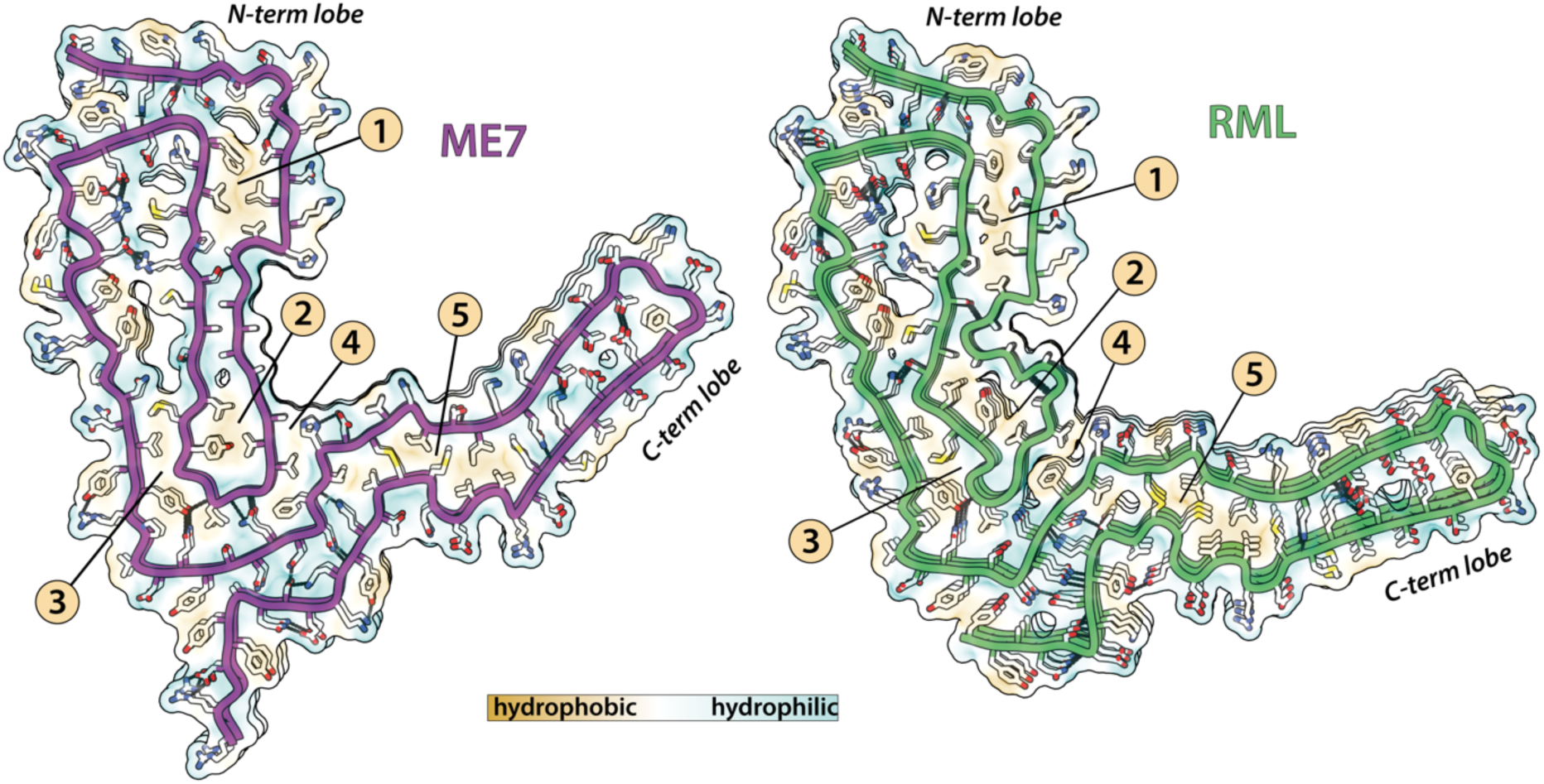
Lateral interactions stabilising mouse prion fibrils. Depiction of polar and non-polar lateral contacts. Protein backbone is shown with cartoon (licorice) representation, amino acid side chains as white sticks coloured by heteroatom (O, red; N, blue; S, yellow) and selected hydrogen bonds as black lines. The hydrophobic patches are numbered 1-5.

**Supplementary Figure 6.**
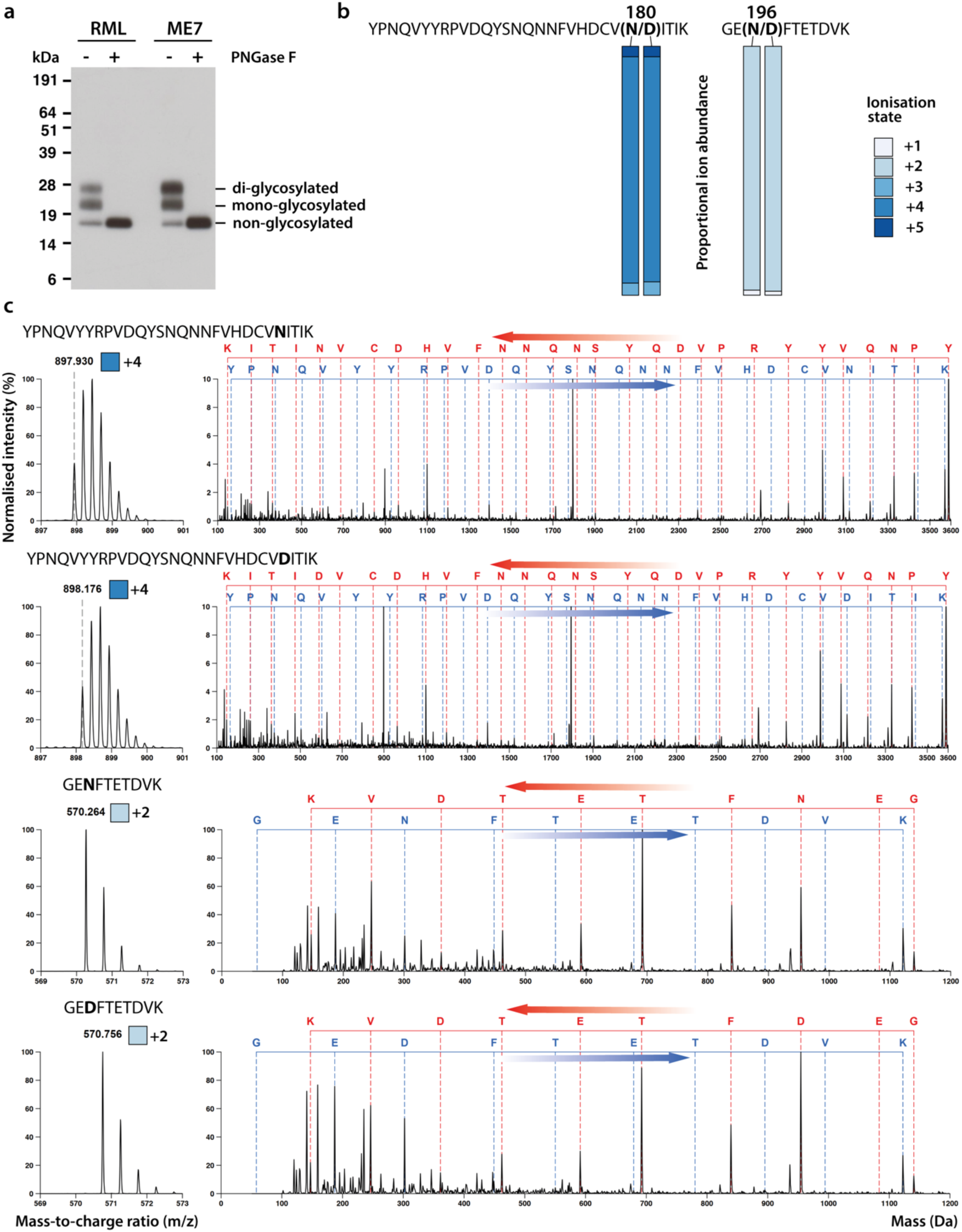
Method validation for determination of glycan occupancy by MS. **a** Western blot confirmation of complete de-glycosylation of PrP in purified *ex vivo* rod preparations by PNGase F. **b** Proportional abundance of all ionisation states found for each peptide reporting N180- and N196-glycan occupancy (bar = 100%); N and D versions show similar distribution. **c** Identification of non- and de-glycosylated peptide ions by MSMS spectral peak assignments of parent ions (left panels) and collision-induced fragmentation spectra (right panels). The most abundant ionisation states of each peptide were used.

**Supplementary Figure 7.**
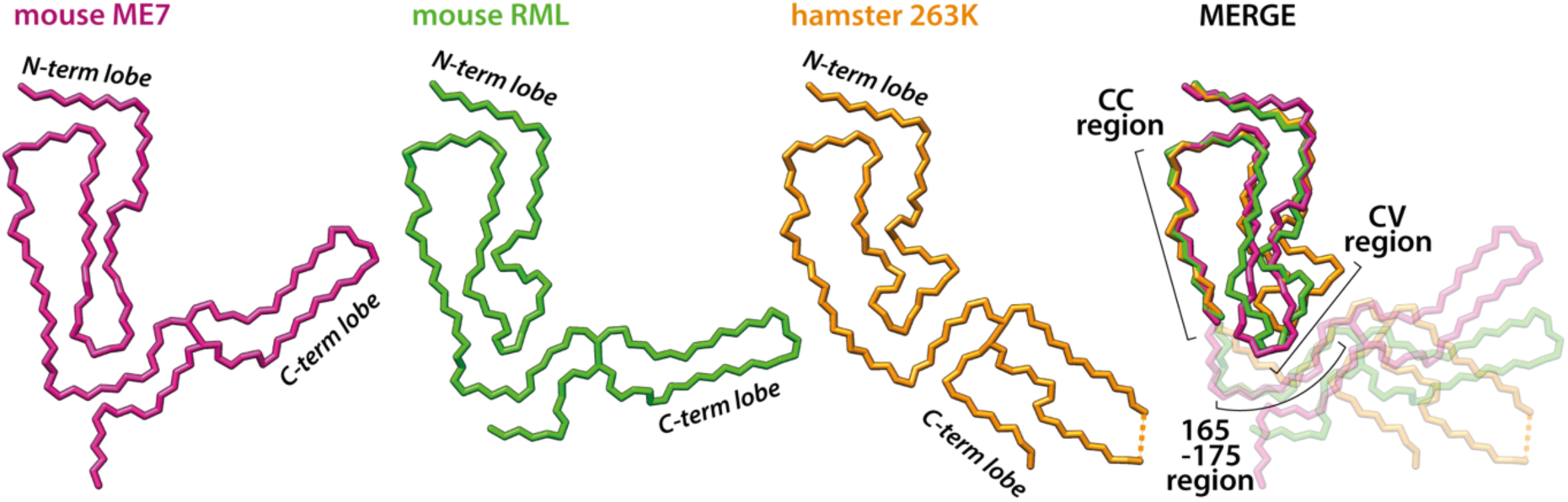
Comparison of rodent prion structures. Backbones of the three structures are aligned on the conformationally conserved (CC) region in the N-terminal lobe (MERGE, dimmed beyond the CC region). The C-terminal lobe of mouse prion strains has divergent orientations due to differences in the N-terminal lobe’s conformationally variable (CV) region that interfaces with the C-terminal lobe. The C-terminal lobe of the hamster strain diverges further from the mouse strains due to additional differences in its primary structure (PrP sequence). The 165-176 region, which corresponds to the β2-α2 loop in PrP^C^, is part of the inter-lobe interface in all three rodent prion fibril structures.

